# Single cell transcriptomics reveals spatial and temporal dynamics of gene expression in the developing mouse spinal cord

**DOI:** 10.1101/472415

**Authors:** Julien Delile, Teresa Rayon, Manuela Melchionda, Amelia Edwards, James Briscoe, Andreas Sagner

## Abstract

The coordinated spatial and temporal regulation of gene expression in the vertebrate neural tube determines the identity of neural progenitors and the function and physiology of the neurons they generate. Progress has been made deciphering the gene regulatory programmes responsible for this process, however, the complexity of the tissue has hampered the systematic analysis of the network and the underlying mechanisms. To address this, we used single cell mRNA sequencing to profile cervical and thoracic regions of the developing mouse neural tube between embryonic days (e)9.5-e13.5. We confirmed the data accurately recapitulates neural tube development, allowing us to identify new markers for specific progenitor and neuronal populations. In addition, the analysis highlighted a previously underappreciated temporal component to the mechanisms generating neuronal diversity and revealed common features in the sequence of transcriptional events that lead to the differentiation of specific neuronal subtypes. Together the data provide a compendium of gene expression for classifying spinal cord cell types that will support future studies of neural tube development, function, and disease.

## INTRODUCTION

Neuronal circuits in the spinal cord receive and process incoming sensory information from the periphery, and control motor output to coordinate movement and locomotion (Goulding, 2009; Kiehn, 2016). The assembly of these circuits begins around embryonic day (e)9 in mouse embryos with the generation of distinct classes of neurons from proliferating progenitor cells located at defined positions within the neural tube. This is directed by signals emanating from the dorsal and ventral poles of the neural tube that partition progenitors into 13 transcriptionally distinct domains ordered along the dorsal ventral axis (Alaynick et al., 2011; Briscoe and Small, 2015; Jessell, 2000; Lai et al., 2016; Le Dréau and Martí, 2012). The gene expression programme of each progenitor domain determines the neuronal cell type it generates (Jessell, 2000; Lee and Pfaff, 2001). Neurons differentiate asynchronously from progenitors by undergoing a series of transcriptional changes that converts a proliferative progenitor into specific classes of neurons. Post-mitotic neurons subsequently further diversify into discrete subsets of physiologically distinct neuronal subtypes and commence formation of the circuitry characteristic of the spinal cord (Bikoff et al., 2016; Borowska et al., 2013; Goulding, 2009; Hayashi et al., 2018; Kiehn, 2016; Lu et al., 2015). Following the period of neurogenesis, which lasts until ∼e13 in mouse, the remaining undifferentiated progenitors in the neural tube produce glia cells. This is accompanied by specific changes in gene expression in progenitors (Deneen et al., 2006; Kang et al., 2012).

Although aspects of the gene regulatory network controlling neural tube patterning and neuronal differentiation have been characterised, our knowledge remains partial and there are still substantial gaps in our understanding. For instance, it is unclear whether the complete catalogue of transcriptional regulators defining cell types has been established. Moreover, whether there are features in common between the mechanisms that promote the differentiation of distinct neuronal subtypes is not known. Similarly, the alterations in gene expression that accompany temporal changes in progenitor competence and cell type generation are poorly documented. In part this lack of knowledge is because, until recently, systematic expression profiling studies have been limited to the use of bulk dissected material. The spatial complexity of the neural tube and the lack of developmental synchrony between cells means that bulk studies provide information on average transcriptional changes but do not have the resolution or sensitivity to provide detailed insight into gene expression or dynamics in specific cell lineages. Conversely, available single cell expression profiling studies of neural development have either focussed on *in vitro* derived neural progenitor cells (Briggs et al., 2017; Sagner et al., 2018), or analysed cells from post-natal animals or late stage embryos (Häring et al., 2018; Rosenberg et al., 2018; Sathyamurthy et al., 2018; Saunders et al., 2018; Zeisel et al., 2018).

To define systematically the complexity of cell types in the developping neural tube and determine the sequence of transcriptional events associated with neurogenesis and changes in progenitor competence, we performed single cell mRNA-Seq analysis. We recovered 21465 cells from cervical and thoracic regions of the mouse neural tube across 5 developmental time points from e9.5 to e13.5. These included cells representing each major spinal cord cell type. The data provide an unbiased classification of neural tube cell populations and their associated gene expression profiles. Using this dataset, we were able to infer the developmental trajectories leading to distinct neural cell fates and to identify cohorts of co-regulated genes involved in specific developmental processes. This compendium of gene expression provides a molecular description of neural tube development and is a resource that suggests testable hypotheses about neural tube development, function, and disease.

## RESULTS

### Assignment of transcriptomes to cell identities

To generate a gene expression atlas of the developing mouse neural tube, we used droplet-based scRNA-seq (10x Genomics Chromium) of microdissected and dissociated cervical and thoracic regions of the spinal cord from mouse embryos between e9.5 and e13.5 (Fig 1A). We generated two replicates per timepoint for e9.5, e10.5 and e11.5. To compensate for the increase in size and cell number of the spinal cord at later developmental stages, three replicates per timepoint were generated for e12.5 and e13.5 (Fig S1A). In total, 41,025 cells were sequenced. After applying quality filters, a dataset of 38976 cells was retained for further analysis (7476 cells at stage e9.5, 6769 cells from stage e10.5, 6634 cells from stage e11.5, 8711 cells from stage e12.5, 9386 cells from e13.5). The average number of UMIs and detected genes in these cells was similar between all samples analysed (Fig S1B,C).

**Figure 1:**
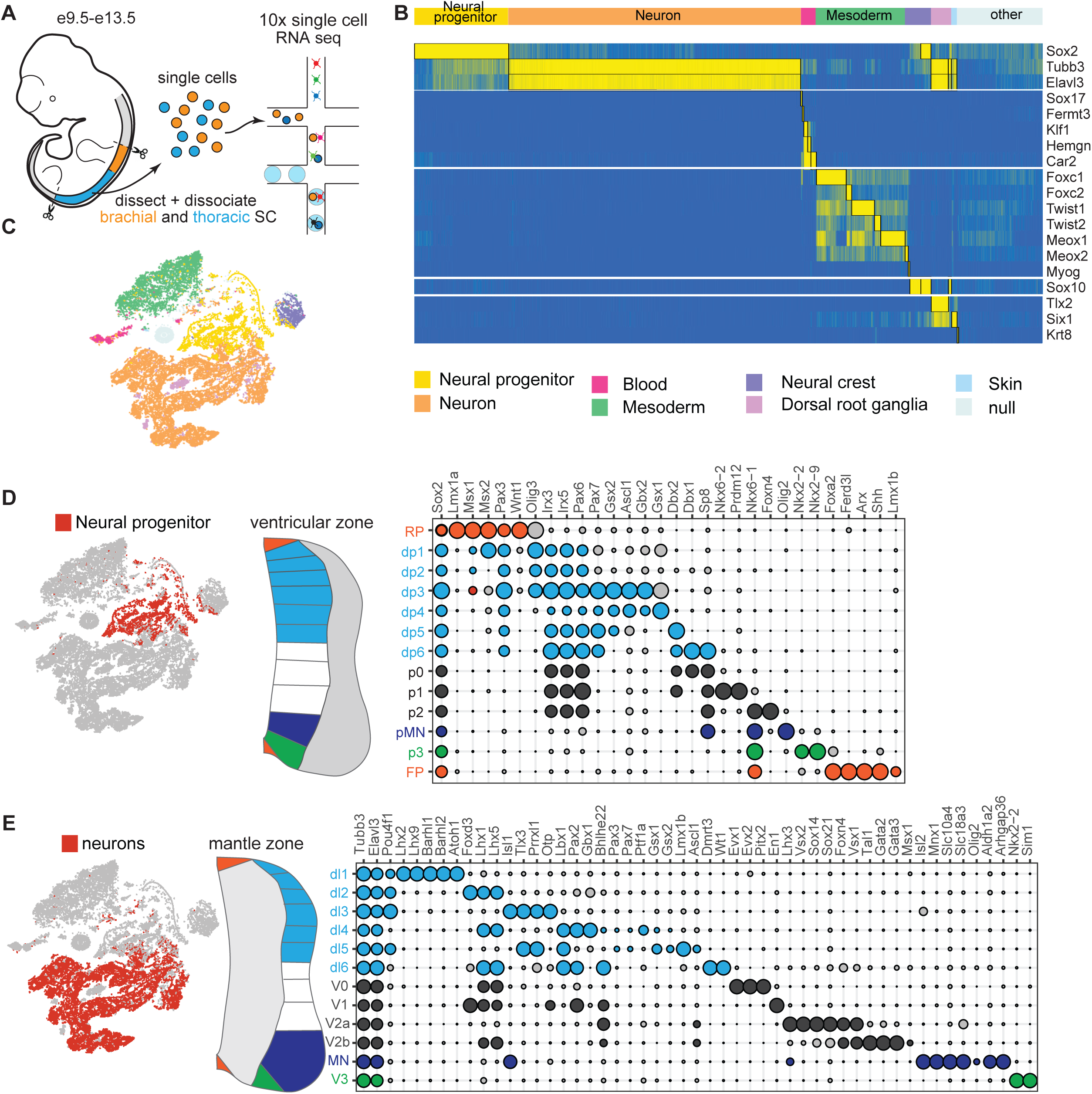
High throughput scRNA-seq from the developing spinal cord. (A) Brachial (orange) and thoracic (blue) regions of the spinal cord of mouse embryos between stage e9.5 to e13.5 were dissected, dissociated and sequenced using the Chromium 10x Genomics system. (B) Partitioning of cells to specific tissue types based on the combinatorial expression of known markers. (C) t-distributed Stochastic Neighbor Embedding (tSNE) plot of the entire dataset based on transcriptional similarity using the same markers as (B), coloured by assigned cell type. Neural progenitors (yellow) and neurons (orange) were selected for further analysis. (D-E) Bubble charts depicting the expression of markers used to identify different dorsal-ventral domains of progenitors (D) and neuronal classes (E). The size of the circles indicates normalized gene expression levels. Genes selected for cell categorisation are coloured; grey circles correspond to markers not used for the selection of a specific population.

To assign identities to the cells in the dataset, we first allocated cells to different tissues based on the combinatorial expression of a curated gene list (Fig 1B). To this end, we used Sox2 to identify spinal cord neural progenitors; Tubb3 and Elavl3 as markers for spinal cord neurons; Fermt3, Klf1, Hemgn and Car2 for blood cells; Foxc1, Foxc2, Twist1, Twist2, Meox1, and Meox2 for early mesodermal cells; Myog1 for myoblasts; Sox10 and Sox2 for neural crest cells; Tubb3, Elavl3, Sox10, Six1 and Tlx2 for neural crest neurons; and Krt8 for skin cells. This allowed the classification of 33754 cells, 87% of the total cells. The remaining cells did not fall into any of the categories, potentially because of poorly resolved transcriptomes or because these cells were derived from other tissues of the embryo. Further subclustering of the unclassified population did not reveal cell populations with spinal cord identity. We also estimated the rate at which the individual transcriptomes might represent more than one cell by assessing the proportion of transcriptomes displaying a gene expression signature of both neural and mesodermal tissue. This indicated that ∼1% of the transcriptomes were from a mixture of neural and mesodermal cell types (Fig. S1D).

Visualizing the resulting dataset with tSNE dimensionality reduction revealed the separation of cells into multiple groupings that reflected the anticipated cell types (Fig 1C). Cells from replicate embryos, whether of the same or different sex, tended to intermingle within the embedding, suggesting minimal batch variation between embryos (Fig S1E,F). By contrast, there were obvious differences in the proportions of cell types at the different timepoints. Most neurons were contained in the datasets obtained at later developmental stages (Figs 1C and S1E), consistent with the proportion of neurons increasing in the spinal cord over time (Kicheva et al., 2014). Conversely, most of the neural crest and mesoderm cells originated from e9.5 and e10.5 embryos (Figs 1C and S1E). This was due to the difficulty of cleanly dissecting the neural tube without accompanying adjacent tissue at these developmental stages.

As it was our aim to construct a spatiotemporal gene expression atlas of the developing spinal cord, we focused on the cells identified as spinal cord neural progenitors and neurons. Extensive characterization of the different populations of neural progenitors and neurons in the spinal cord has revealed a variety of molecular markers that specifically or combinatorially define different domains of progenitors and classes of neurons (Alaynick et al., 2011; Lai et al., 2016). Taking advantage of this, we first generated a binary ‘knowledge matrix’ in which each progenitor domain and neuronal class is defined by the expression of a characteristic marker set (see Table S1). Notably, a specific cell type can be defined by more than one combination of marker gene expression patterns, which helps to capture certain subpopulations, such as early neurons that still partially express progenitor markers and have not yet fully activated their neuronal gene expression programmes. We then binarized the expression profiles of the defined marker genes in the transcriptome data. Cells in the dataset were then assigned an identity from the knowledge matrix, using the minimal Euclidean distance between each transcriptome and cell type classes. Plotting the resulting average levels of marker gene expression for the assigned cell identities revealed the well-known patterns of progenitor- and neuron-specific gene expression patterns (Fig 1D-E and S1F). We therefore conclude that the partitioning algorithm correctly assigns identities to the cells in our dataset.

### Developmental dynamics of cell types

We hypothesized that sampling several thousand transcriptomes per time point would be sufficient to reconstruct the changes in domain sizes during development. Plotting the proportion of progenitors and neurons over time revealed a sharp decrease in the overall proportion of progenitors from >65% of cells at e9.5 to <15% of cells at e13.5 (Fig. 2A). Previous work suggested that progenitor domain sizes in the spinal cord are dictated by two processes (Kicheva et al., 2014). At early stages morphogens emanating from the poles establish the general bauplan of the spinal cord by inducing distinct progenitor identities along the dorsal-ventral axis, while at later stages the relative size of these progenitor domains is dictated by domain-specific rates of neuronal differentiation. To test if the proportions of cells recovered from the assignment of cell identities correctly recapitulated the growth dynamics of the spinal cord, we compared with the quantification in Kicheva et al (2014). To this end we classified cells in our dataset as FP, p3, motor neuron (MN) progenitors (pMN), and the broader domains pI (composed of p0, p1 and p2) and pD (including pd1, pd2, pd3, pd4, pd5, pd6) (Fig. 2E). This revealed a close match in the dynamics predicted by the two independent datasets. The correspondence between the data allowed us to extend the analysis beyond the E11.5 end point of Kicheva et al (2014). Between e11.5 and e12.5, the rate of neurogenesis in the spinal cord appears to slow, leading to a stabilization of the proportion of progenitors in all domains (Fig 2B,E); 24h later, at e13.5, the fraction of progenitors in dorsal domains decreased relative to ventral (Fig. 2B,E). This is in line with the previously observed higher rate of neurogenesis in the dorsal spinal cord at e13.5 and the consequent depletion of progenitors (Gross et al., 2002; Lai et al., 2016; Müller et al., 2002).

**Figure 2:**
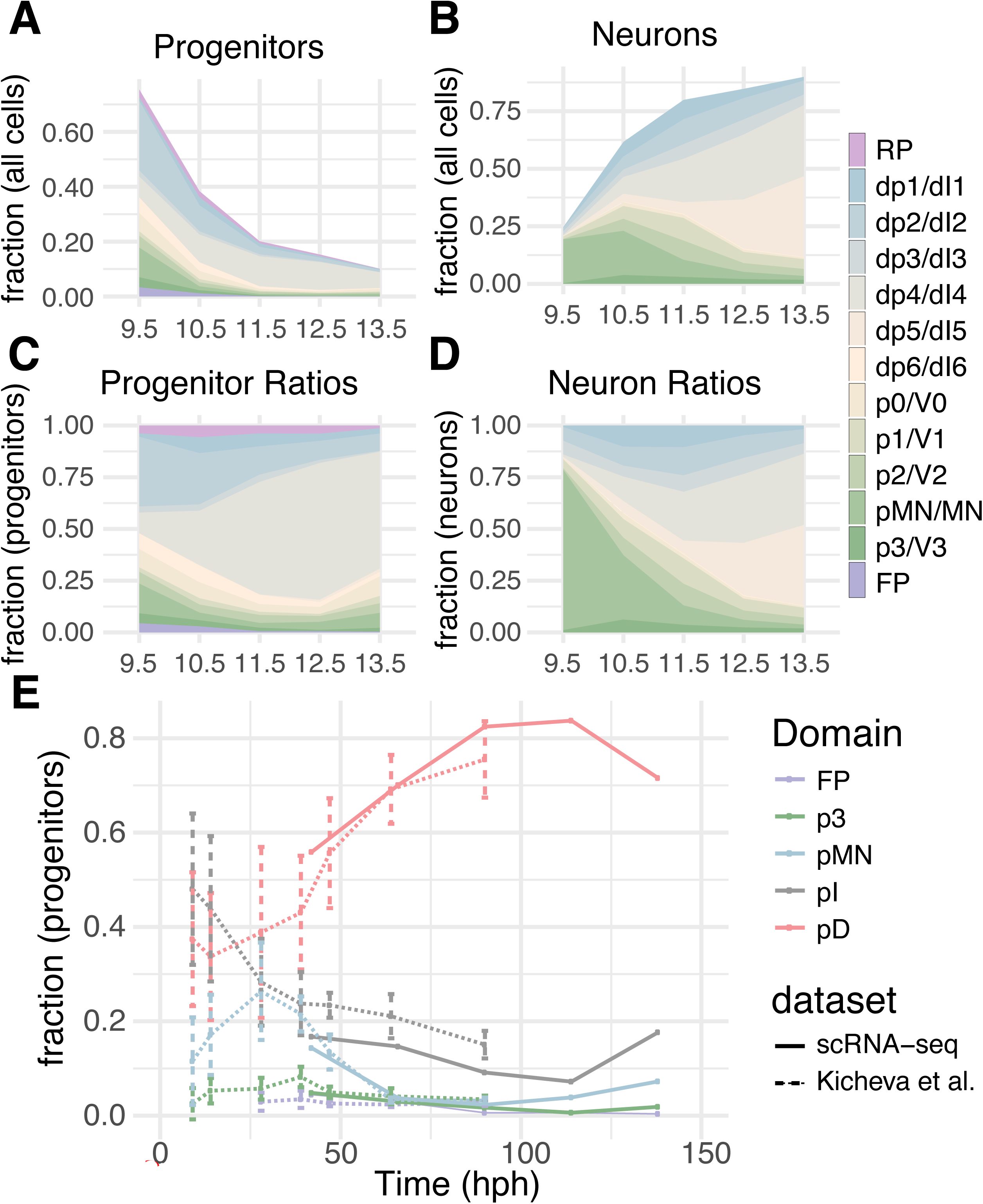
Dynamics of domain sizes based on the single cell sequencing data. (A-D) Changes in the fraction of progenitors (A,C) and neurons (B,D) between e9.5 and e13.5. (A,B) The data are normalised to the sum of neurons and progenitors detected at each time point. (C,D) show fractions within progenitors (B) or neurons (D). (E) Comparison of the ratio of progenitors from our dataset (solid lines) with those of Kicheva et al. (2014; dashed lines). The broader domains pI and pD are composed of p0-p2 and pd1-pd6 progenitors, respectively.

We performed a similar analysis on the population dynamics of the neuron subtypes (Fig. 2C,D). The most prominent class of neurons at early developmental stages (e9.5 and e10.5) are MNs, consistent with the higher initial differentiation rate of MN progenitors compared to other neural progenitors in the spinal cord (Ericson et al., 1992; Kicheva et al., 2014; Novitch et al., 2001; Sagner et al., 2018) (Fig. 2C,D). At later developmental stages (after e11.5) the proportions of inhibitory dI4 or excitatory dI5 neurons increased markedly (Fig. 2D). This is explained by the expansion of the combined dp4/dp5 progenitor pool in the dorsal spinal cord at these stages and the extended phase of neurogenesis in the dorsal spinal cord that results in the formation of late born dI4 and dI5 neurons (also known as dILa and dILb) (Gross et al., 2002; Müller et al., 2002). Based on these observations, we conclude that the single cell transcriptomic atlas correctly captures the domain dynamics of progenitor and neuronal populations.

### Prediction of novel gene expression patterns

We next sought to extend the analysis by identifying further cell type specific expression patterns. Here the challenge is that different cell types are defined by the intersection of overlapping combinations of gene expression. These combinations do not necessarily identify adjacent cell types. To overcome this, we constructed the list of all 2^13^ models of combinatorial patterns in progenitor domains (8192 combinations from 13 progenitor domains, including floor plate and roof plate) that we used to identify genes with differential expression patterns. For each gene, the best fit model was selected and, after filtering by fold-change and significance level, we obtained a list of 102 categories of combinations (Fig. S2). The same procedure was then applied to the 12 neuronal populations (V3, MN, V2a, V2b, V1, V0, dl6, dl5, dl4, dl3, dl2, dl1). From the 4096 combinatorial models obtained, we predicted 147 categories of distinct patterns. We applied a conservative p-value cutoff of 10^-9^ to obtain a manageable list of candidates from the differential gene expression analysis; lowering this cutoff will extend the list of candidates to pursue in the future.

For further analysis, we first focused on genes encoding proteins involved in neurotransmitter biogenesis and release (Fig. 3A). These identified the three main types of spinal cord neurons: cholinergic MNs, inhibitory GABAergic and glycinergic interneurons, and excitatory glutamatergic interneurons. As expected, these could be distinguished based on the expression of metabolic enzymes necessary for neurotransmitter production (e.g. Glutamatedecarboxylases Gad1 and Gad2 necessary for transmitter production) and specific vesicular transporters e.g. Slc18a3 and Slc5a7 (vAcht and high-affinity choline transporter in MNs), Slc32a1 and Slc6a5 (vIAAT and Glycine Transporter 2 [GlyT2] in inhibitory interneurons) and Slc17a6 (vGlut2 in excitatory neurons). The analysis correctly predicted expression in the different neuronal populations: cholinergic MNs, inhibitory GABAergic and glycinergic dI4, V1 and V2b interneurons and excitatory glutamatergic dI1, dI2, dI3, dI5, V2a, and V3 interneurons (Alaynick et al., 2011; Lai et al., 2016; Lu et al., 2015).

**Figure 3:**
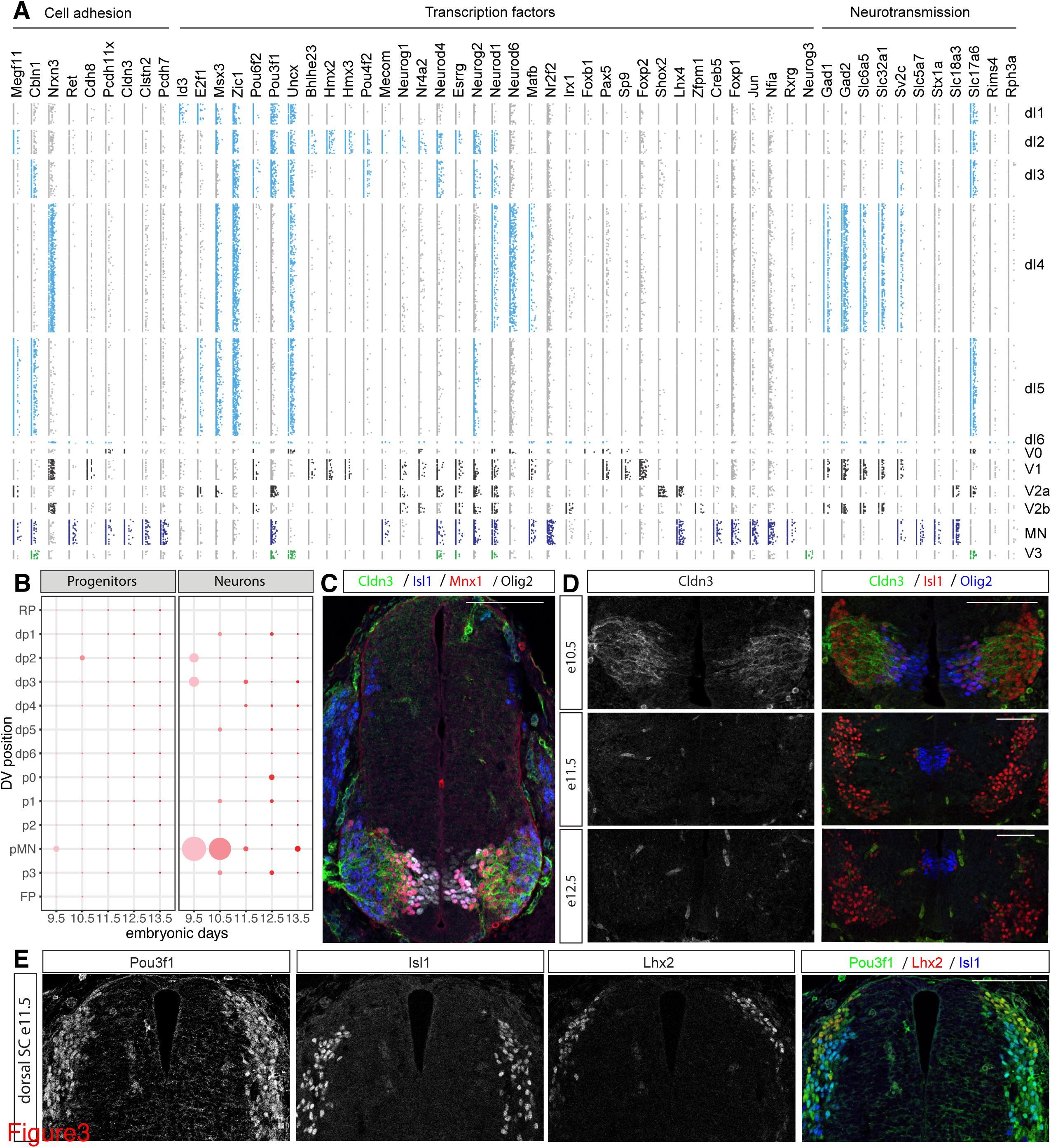
Spatial and temporal patterns of gene expression in neural progenitors and neurons. (A) Identification of differentially expressed genes encoding cell adhesion molecules, TFs and proteins involved in neurotransmission. Predicted genes that were used in the initial partitioning of cell types are not shown. (B) Spatial and temporal expression of Cldn3 in neural progenitors and neurons in the dataset. Cldn3 is specifically expressed in MNs until e10.5. (C) Immunostaining at e10.5 for Cldn3 (green), Isl1 (blue), Mnx1 (red) and Olig2 (grey). (D) Cldn3 expression is specific to MNs at e10.5, and expression is lost in MNs at e11.5 and e12.5. (E) Pou3f1 is expressed in dorsal d1-3 neurons at e11.5. dI1 neurons are labelled by Lhx2, and dI3 neurons by Isl1 expression. Scale bars = 100 µm

We next examined adhesion molecules (Fig 3A). In the spinal cord, differential cell adhesion is essential for the clustering of MNs into distinct pools (Demireva et al., 2011; Price et al., 2002). Furthermore, differential adhesion mediated by Neurexins and Cerebellins is required for synapse formation and function (Südhof, 2017; Uemura et al., 2010). Interrogation of the transcriptome data recovered 9 differentially expressed cell adhesion molecules (Cldn3, Cbln1, Cdh8, Pcdh11x, Clstn2, Pcdh7, Megf11, Nrxn3, Ret). Consistent with previous work, the analysis correctly predicted Pcdh7, Megf11 and Ret to be expressed in MNs (Catela et al., 2016; Hanley et al., 2016; Lin et al., 2012). Furthermore, we identified Nrxn3 to be specific for inhibitory neurons in the dI4, dI6, V1 and V2b domains. This pattern was anticorrelated with the expression pattern of Cbln1, which was specifically expressed in excitatory dI3, dI5 and V3 neurons and MNs. Moreover, further investigation of other cell adhesion proteins identified Nrxn1, which was slightly below the threshold we set for gene level fold-change in the transcriptome-wide analysis, as specifically expressed in excitatory neurons (dl1, dl2, dl3, dl5) (data not shown). This raises the possibility of a molecular adhesion code that distinguishes inhibitory and excitatory interneurons in the spinal cord, which might contribute to synaptogenesis during circuit formation.

The availability of transcriptome data from multiple time points highlighted the dynamics of gene expression in different cell types. To test whether the dataset could be used to correctly infer spatial and temporal gene expression patterns, we focussed on Cldn3. Cldn3 encodes for a tight junction component (Morita et al., 1999) and was predicted to be specifically expressed in MNs before, but not after, e11.5 (Fig 3B). We performed immunofluorescence assays for Cldn3 on transverse spinal cord sections from e10.5-e12.5 embryos. As predicted by the bioinformatic analysis, Cldn3 expression at e10.5 was specific to MNs and expression was not detectable in e11.5 and e12.5 sections (Fig 3C,D).

We next turned our attention to transcription factors (TFs) and searched for TFs that were differentially expressed between neuronal subtypes (Fig 3A). This recovered multiple TFs, many with well-established differential gene expression patterns, for example: Shox2 expression in V2a neurons (Dougherty et al., 2013; Hayashi et al., 2018); Foxp2 expression in V1 neurons (Morikawa et al., 2009; Stam et al., 2012); Lhx4 expression in MNs and V2a neurons (Lee and Pfaff, 2001; Sharma et al., 1998); expression of Neurog3 and Uncx in V3 neurons (Carcagno et al., 2014; Sommer et al., 1996); and high level expression of Nr2f2 (also known as COUP-TF2), Nfia, Foxp1 and Creb5 in MNs (Dasen et al., 2008; Glasgow et al., 2017; Hanley et al., 2016; Lutz et al., 1994; Rousso et al., 2008). This analysis also suggested multiple previously unknown expression patterns of TFs, for example: expression of Hmx2 and Hmx3 in V1 and dI2 neurons; Sp9 in V1 neurons; and Bhlhe23 in dI2 neurons (Fig 3A).

For validation, we focussed on Pou3f1. This TF has been previously demonstrated to be expressed in multiple neuronal subtypes of spinal cord, e.g in MNs, V2a and V3 interneurons (Dasen et al., 2005; Francius et al., 2013), which we correctly identified. Our analysis also suggested that Pou3f1 is expressed in dorsal dI1-dI3 neurons (Fig 3A), which to our knowledge had not been described before. To test this prediction, we assayed sections of embryonic spinal cord at e11.5 for Pou3f1, the dI1 marker Lhx2, and the dI3 marker Isl1. This analysis revealed broad expression of Pou3f1 in dorsal spinal cord neurons and colocalization between Pou3f1, Lhx2 and Isl1 (Fig 3E). However, we also detected Pou3f1+ cells, which were not expressing Lhx2 nor Isl1, these are presumably dI2 neurons. Thus, the differential gene expression analysis correctly identified the restricted expression patterns of multiple known genes and predicted novel patterns of expression. The resource may therefore facilitate further investigation of the dynamics of gene expression and the gene regulatory network that underlies neuronal differentiation in the spinal cord.

### Clustering of neurons predicts novel neuronal subtypes and transcriptional codes

The 11 neural progenitor domains (p0-p3, pMN, pd1-pd6) generate distinct classes of spinal neurons that further diversify into multiple subpopulations (Alaynick et al., 2011; Bikoff et al., 2016; Du et al., 2015; Francius et al., 2013; Hayashi et al., 2018; Lai et al., 2016; Sweeney et al., 2018). Gene expression profiles that distinguish subpopulations of neurons are well documented. To further probe the resolution of our atlas and identify novel components of the gene regulatory networks that might mediate neuronal subtype specification, we performed hierarchical clustering independently on each neuronal class (Fig 4A-D and Fig S4). To this end, we first identified gene modules consisting of genes demonstrating concerted pattern of expression in each neuron class. The modules that showed differential expression in a class were used to perform hierarchical clustering and assign distinct subtype identities. This resulted in the identification of 59 subpopulations (Fig 4A). Of these, initial inspection indicated only two subpopulations were incorrectly assigned. The first group consisted of poorly characterized neurons lacking any salient transcriptomic feature (clade MN.5) and the second a population expressing late neural progenitor markers Sox9 and Fabp7 indicative of progenitor identity (clade V2a.5).

**Figure 4:**
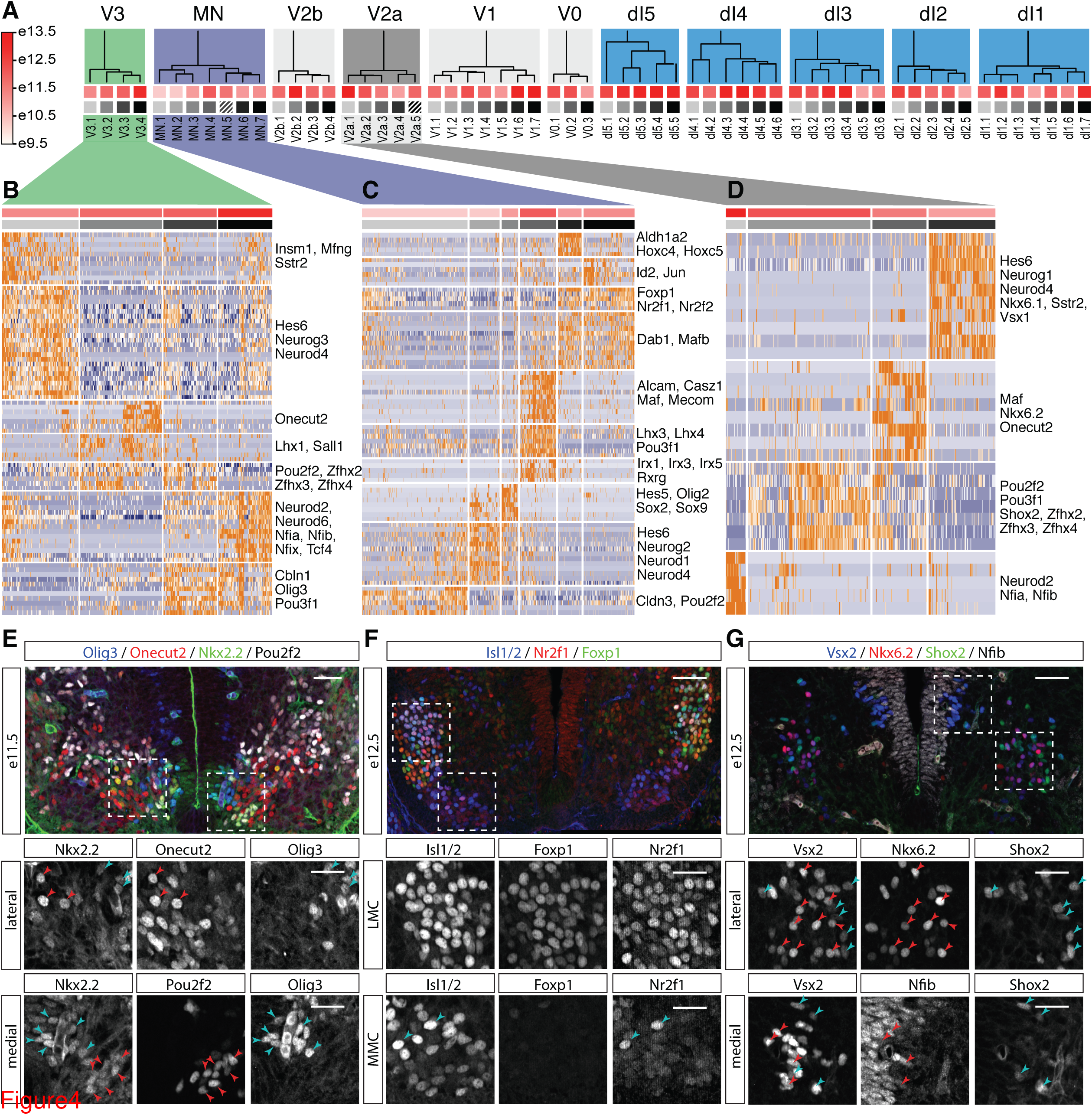
Hierarchical clustering identifies neuronal subtypes and implicates TFs in determining their identity. (A) Hierarchical clustering of the cardinal types of spinal cord neurons reveals 59 neuronal subtypes. Dendrograms for each neuronal domain are depicted. Squares under the dendrogram indicate average age (red squares) and neuronal subtype identity (grey squares). Colours of squares correspond to those shown on top of the heatmaps in Fig. 2B-D and Fig S2. Clades associated with striped squares correspond to incorrectly classified cells that were discarded from further analysis. (B-D) Identification of neuronal subtypes by clustering of gene expression profiles in V3 (B), MNs (C) and V2a interneurons (D). Hierarchical clustering was performed using the indicated gene modules (Table S2). A subset of genes included in the modules is indicated on the righthand side. (E-G) Validation of predicted gene expression patterns obtained from the hierarchical clustering in (B-D). (E) Expression of Pou2f2 is detected in Nkx2.2-expressing V3 neurons at e11.5. Pou2f2 expression does not overlap with Onecut2 in more dorsal V3 neurons (top row) or Olig3 in V3 neurons abutting the p3 domain (bottom row). (F) Nr2f1 expression in Foxp1+ LMC neurons at e12.5 (top row). A few Nr2f1-positive cells are also detected in Foxp1-negative MNs within the MMC (bottom row). (G) Nkx6.2 expression in lateral Vsx2-expressing V2a neurons does not overlap Shox2 expression at e12.5 (top row). Nfib expression is confined to medial V2a neurons at this stage (bottom row). Scale bars = 50 µm, scale bars of insets are 25 µm

To explore whether the approach correctly classified multiple neuronal subpopulations and allowed the identification of novel TFs implicated in the specification of distinct subtype identities, we focused on the identification of subpopulations of neurons in three ventral domains (V2a and V3 interneurons, and MNs) (Fig 4B-D).

V3 interneurons have been previously shown to be comprised of at least 2 subtypes with distinct properties and settling positions in the spinal cord (Borowska et al., 2013; Francius et al., 2013; Goulding, 2009; Lu et al., 2015). In e12.5 mice, these 2 subpopulations can be distinguished molecularly, with dorsal V3d neurons expressing Onecut TFs, while ventral V3v neurons express Olig3 (Francius et al., 2013). Hierarchical clustering of V3 neurons revealed at least 4 different subtypes (Figure 4B). Clade V3.1 consisted of newly differentiating V3 neurons that expressed the neurogenic markers Neurog3, Hes6 and Neurod4. Clade V3.2 was characterized by expression of Lhx1 and could be further subdivided into a Onecut2 positive and negative population. Clade V3.3 expressed the TFs Pou2f2, Zfhx2, Zfhx3, and Zfhx4, while cells in clade V3.4 expressed Neurod2, Neurod6 and Nfia/b/x. Furthermore, clades V3.3 and V3.4 are characterized by the expression of Pou3f1 and Olig3. Thus, the data correctly identified the different known types of V3 interneurons and predicted the existence of an additional Nfia/b/x and Neurod2/6 expressing subtype (Fig 4B).

After their generation, MNs diversify and organize into distinct columns and pools along the anterior-posterior axis of the spinal cord (Francius and Clotman, 2014; Jessell, 2000; Philippidou and Dasen, 2013; Stifani, 2014). Subclustering of the identified MNs revealed 6 clades (Figure 4A): Clades MN.2 and MN.3 comprised neurogenic progenitors because cells in these clades expressed the progenitor markers Sox2, Hes5 and Olig2 (clade MN.3), or Hes6, Neurog2, and Neurod4 (clade MN.2) and did not express substantial levels of the vesicular acetylcholine transporter Slc18a3. The remaining four clades corresponded to more mature neurons. The remaining four corresponded to early differentiated MNs: clade MN.1 characterized by the expression of the TF Pou2f2 and the tight junction component Cldn3; Median Motor Column (MMC) neurons and Phrenic Motor Column (PMC) neurons grouped in clade MN.4, which had high levels of Lhx3, Mecom, Pou3f1 (also known as Oct6 or SCIP) and the PMC marker Alcam (Hanley et al., 2016; Stifani, 2014); and 2 clades (MN.6 and MN.7) of Foxp1-positive cells were identified. The Foxp1 expressing MNs correspond to cervical Lateral Motor Column (LMC) neurons (clade MN.6) and thoracic preganglionic motor column (PGC) neurons (clade MN.7). Clade MN.6 was characterized by the expression of a gene module containing Aldh1a2 (also known as Raldh2) and the cervical Hox genes Hoxc4 and Hoxc5 (Fig 4C). Furthermore, cells in this clade did not express Hoxc9, consistent with their limb level location of the LMC. By contrast, cells in clade MN.7 expressed higher levels of Hoxc9, characteristic of the thoracic regions (data not shown).

V2a interneurons have recently been shown to consist of two major subgroups that have different settling positions in the neural tube: a medial subgroup expressing Nfib and Neurod2 and a lateral subgroup that expresses Shox2 and Zfhx3 (Hayashi et al., 2018). Furthermore, the lateral subgroup seems to further diversify into subgroups labelled by Shox2 and Maf/Onecut TFs (Francius et al., 2013; Harris et al., 2018). Consistent with this, hierarchical clustering of the single cell transcriptome data revealed four major subtypes of V2a neurons (Fig 4D). Clade V2a.4 consisted mainly of newly differentiating neurons that expressed the neurogenic V2a markers Neurog1, Vsx1 and Hes6, but were not expressing the mature V2a marker Vsx2 (also known as Chx10). The three other neuronal subtypes were characterized by the expression of Neurod2, Nfia and Nfib (clade V2a.1), Shox2, Pou3f1, Pou2f2, Zfhx2, Zfhx3 and Zfhx4 (clade V2a.2) and Maf and Onecut2 (clade V2a.3). Thus, the single cell transcriptome data correctly recovers the different subtypes of V2a interneurons.

To test the accuracy of the transcriptome data and our computational assignment of transcriptomes to cell identities we experimentally assayed three predictions: the expression of Pou2f2 in V3, Nr2f1 (also known as COUP-TF1) in MNs and Nkx6.2 in V2a interneurons (Fig 4B-D). Our analysis suggested that these TFs are only expressed in specific subsets of neurons: Pou2f2 in a group of V3 interneurons that is complementary to the Olig3 and Onecut2 expressing populations; Nr2f1 in Foxp1+ MNs; and Nkx6.2 in a subpopulation of V2a interneurons that does not overlap with Shox2 and Nfib. Assaying expression in e11.5 and e12.5 cervical spinal cord (Figure 4E-G) confirmed each of the predictions. We therefore conclude that our spinal cord atlas provides sufficient resolution to detect distinct neuronal subtypes and implicates novel TFs in their specification.

### Modular and temporal specification of neuronal identity

Having compiled a gene expression atlas encompassing multiple cell types and timepoints, we asked if we could use it to investigate the timing of neuronal subtype generation. We first asked whether we could identify sets of TFs that are used recurrently to define subpopulations of neurons in multiple domains. Focussing on the ventral domains (V0-V3 and MNs), we found that neurons in all these classes could be further partitioned using the expression of specific gene modules (Fig. 5A). Modules containing Pou2f2 were expressed in subpopulations of each neuronal class. Furthermore, Pou3f1, Nfib, Onecut and Maf also demarcated subpopulations of multiple neuronal classes, consistent with the expression of these TFs in multiple types of neurons (Francius et al., 2013). The composition of the gene modules containing these TFs was well conserved between neuronal domains. The Pou2f2 gene module also contained Zfhx2, Zfhx3, and Zfhx4, in most cases. By contrast, the Nfib gene module often included Nfia, Nfix, Neurod2, Neurod6 and Tcf4. This raised the possibility that similar transcriptional programs mediate neuronal diversification in each of these domains.

**Figure 5:**
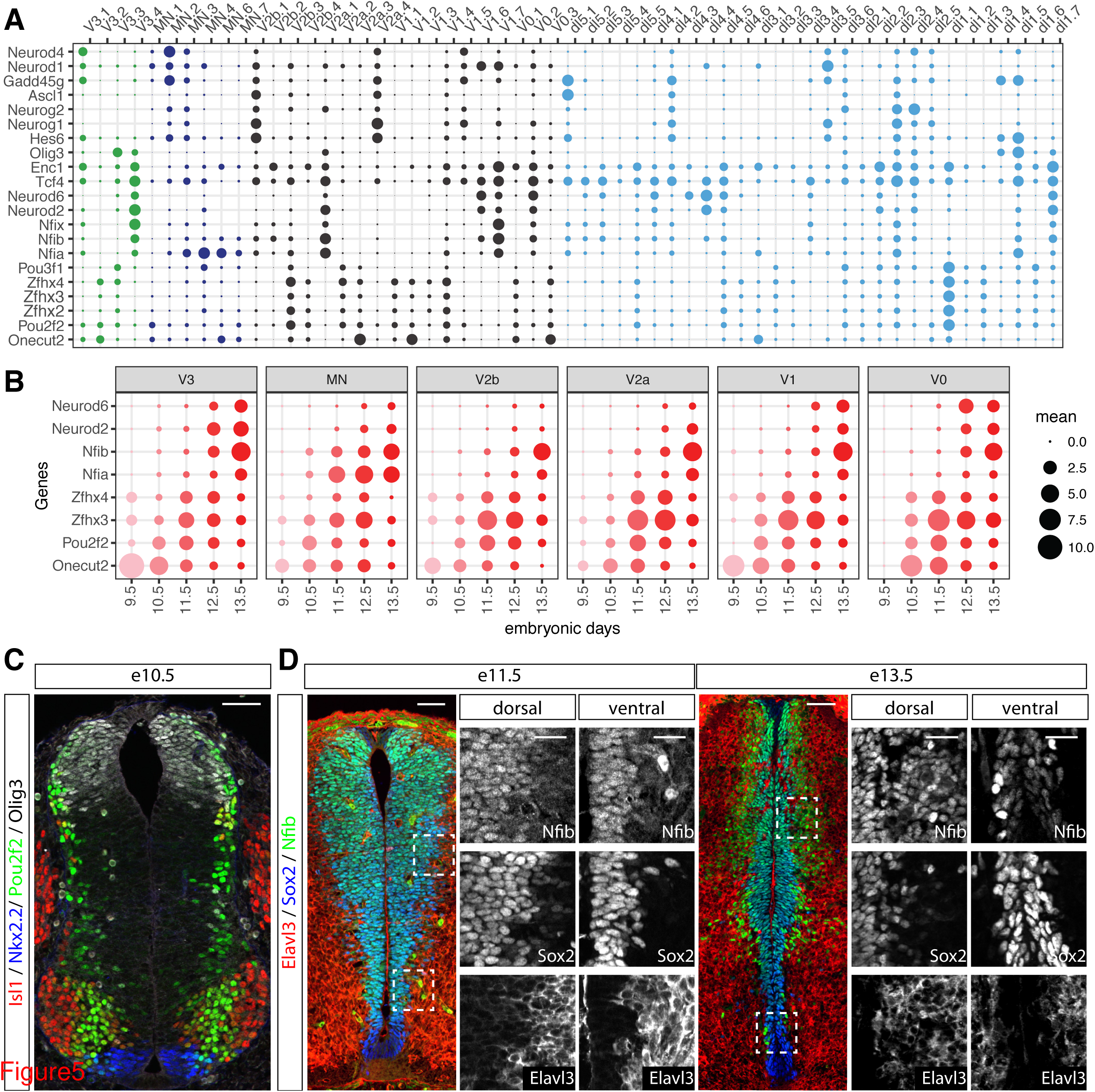
Temporal stratification of neuronal subtypes by shared sets of TFs. (A) Gene expression profiles of TFs that are used repeatedly to define subpopulations of neurons in multiple domains. (B) Temporal expression of a shared set of TFs in different neuronal populations identifies two waves of neurogenesis. The size of the circles indicates the mean expression of genes per stage and domain. (C) Widespread expression of Pou2f2 at early timepoints (e10.5) in differentiating neurons close to the ventricular zone. Pou2f2 expression colocalizes with Olig3, Nkx2.2 and Isl1. (D) Increased expression of Nfib in differentiating neurons at late developmental stages. Nfib is expressed at low levels at e11.5 in progenitors labelled with Sox2 and is not detected in neurons. By contrast, at e13.5 Nfib expression is observed in neurons that also express the neuronal marker Elavl3. Scale bars = 50 µm, scale bars of insets are 25 µm

We asked whether specific gene modules were induced at different timepoints in development. To this end, we focussed on the TFs Onecut2, Pou2f2, Nfib and Neurod2 and assessed the timing of their induction. Plotting the average expression level of these TFs for each neuronal class between e9.5 and e13.5 revealed a temporal stratification of neuronal subtypes that was conserved between domains (Fig. 5B). The proportion of neurons that expressed Onecut2 and Pou2f2 peaked at early developmental stages (before e11.5), while Nfib and Neurod2 expression were induced at e12.5 or e13.5. This is consistent with prior observations. Onecut TFs have been shown to be expressed early in V1 and MNs (Roy et al., 2012; Stam et al., 2012) and Nfia/b are induced at later time points (Deneen et al., 2006; Kang et al., 2012). *In vivo* assays of Pou2f2 confirmed its expression in neurons at e10.5 (Fig 5C), whereas Nfib expression was not expressed in neurons until after e11.5 (Fig 5D). Based on these observations, we conclude that neuronal subtype diversification in the spinal cord is at least partly driven by transcriptional programmes that are shared between multiple domains and sequentially induced in each of the domains. This suggests a temporal component to the specification of neuronal subtype identity that complements the well described spatial axis of patterning.

### Reconstruction of the gene expression dynamics underlying neurogenesis

Finally we turned out attention to the changes in gene expression that accompany the transition of progenitors to neurons. Single cell transcriptome analysis enables the reconstruction of gene expression dynamics along differentiation trajectories (Setty et al., 2016; Shin et al., 2015; Trapnell et al., 2014). To this end, we projected the cells into a 2-dimensional space (Fig 6A) using a Principal Component Analysis from a 100-dimensional space defined by a set of genes expressed in all DV domains during progenitor maturation and neurogenesis (Fig. S6A). The first principal component aligned with neurogenesis, indicated by the down-regulation of Sox2 and up-regulation of Tubb3, hence we used the PC1 coordinates as a pseudo-temporal ordering for neurogenesis. We then reconstructed, independently for each domain, a smoothed expression profile of genes along pseudotime by fitting spline curves (Fig. 6B). This predicted the expression changes in a set of 36 regionally patterned genes involved in neurogenesis (Fig. 6C).

**Figure 6:**
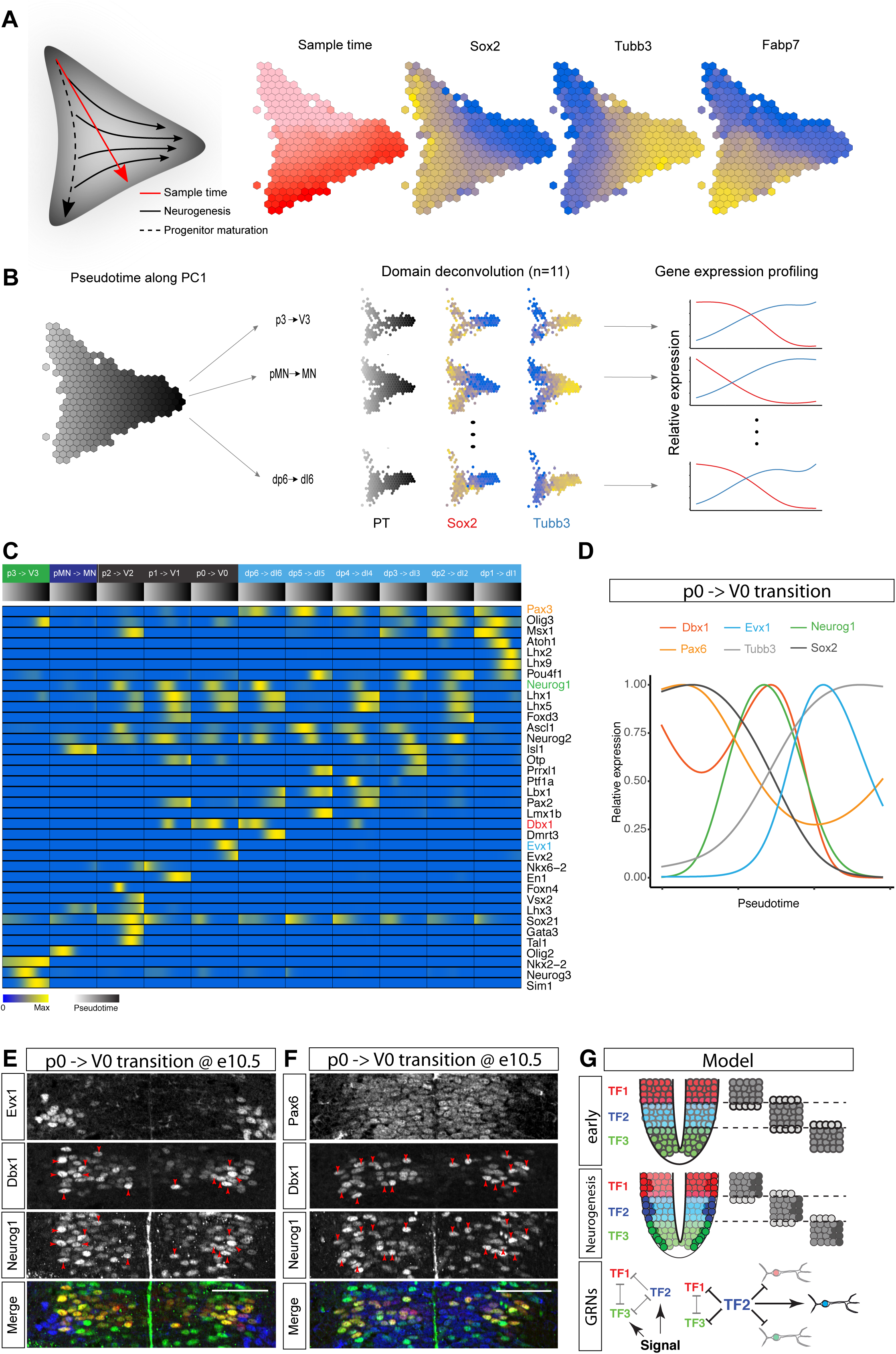
Pseudotemporal ordering reveals gene expression dynamics during neurogenesis. (A) Two-dimensional PCA projection of all neural cells, shown on a hexagonal heatmap, from a 100- dimensional space defined by genes expressed in all DV domains during progenitor maturation and neurogenesis. The graph on the left depicts from top to bottom progenitor maturation, while the arrows from left to right mark neurogenesis. The hexagonal heatmaps indicate the number of cells from different developmental stages, and the expression pattern of the pan-progenitor marker Sox2, the neuronal marker Tubb3, and the gliogenic marker Fabp7. (B) The first principal component of the cell state graph was used to independently reconstruct neurogenesis in each dorso-ventral domain. Cells allocated to specific DV domains were plotted along the differentiation trajectory, and the expression profile of genes was independently reconstructed. (C) Upregulation of domain specific TFs coincides with neurogenesis in multiple domains. Heatmap including the normalized expression pattern (low blue, high yellow) of genes involved in neurogenesis per domain along the pseudotemporal axis (grey early, black late). (D) Smoothed expression profile of the neurogenic trajectory from p0 to V0 shows a transient upregulation of Dbx1 prior to neurogenesis that coincides with the maximal expression of Neurog1. (E-F) Upregulation of Dbx1 coincides with Neurog1 expression. (E) Co-expression of Dbx1 and Neurog1 in cells in the differentiation zone of the ventricular layer in the p0 domain at eE10.5. V0 neurons are identified by the expression of Evx1. (F) While Dbx1 expression is maximal in differentiating progenitors at e10.5, Pax6 expression is homogeneous in all p0 progenitors. (G) Domain-specific TFs are upregulated prior to neurogenesis. Initially, domain-specific TFs set the specificity of the progenitors. Upon neurogenesis, expression is upregulated to reinforce the subtype identity of the differentiating neurons.

Examining gene expression changes as progenitors transitioned to neurons revealed the transient upregulation of several TFs. These included several known neurogenic factors, including Atoh1, Neurog1 and Neurog2, and several domain specific TFs. In line with this, we recently demonstrated upregulation of Olig2 coincides with Neurog2 expression and neurogenesis in MN progenitors (Sagner et al., 2018). To test whether similar expression dynamics occur during neuronal differentiation in other progenitor domains, we first focussed on Olig3, which is expressed in the three most dorsal progenitor domains, dp1-dp3 (Müller et al., 2005). Differentiation of these progenitors into dI1-dI3 neurons depends on the expression of the proneural bHLH proteins: Atoh1 in dp1; Neurog1 in dp2 and Neurog2 in dp2 and dp3 progenitors (Lai et al., 2016). Plotting Olig3 expression dynamics along the differentiation trajectory from dp2 to dI2 neurons or dp3 to dI3 neurons revealed that the maximal expression of Olig3 coincided with the expression of Neurog1 and Neurog2 (Fig. S6C,D). To test the specificity of the transient upregulation of Olig3, we examined the dynamics of Msx1 and the more broadly expressed TF Pax3. Expression of both genes decreased upon differentiation, thus showing different expression dynamics than Olig3 (Fig. S6C,D). Assays of spinal cord sections indeed revealed heterogeneity in Olig3 levels, with higher expression of Olig3 correlating with Neurog1 and Neurog2 expression and low levels of Msx1 (Fig S6F-H). These observations confirm that Olig3 expression is upregulated at the onset of neurogenesis and validated the predictions made by the pseudotemporal ordering.

Similar reconstruction of the p0 to V0 transition predicted that Dbx1 levels should increase during neurogenesis (Fig. 6D). Moreover, levels of the p3 marker Nkx2.2 were predicted to increase at the onset of neurogenesis (Fig. S6B). Assays confirmed these predictions: Dbx1 levels were markedly elevated in Neurog1-positive p0 progenitors (Fig. 6E), while Nkx2.2 levels were increased immediately adjacent to the p3 progenitor domain in Olig3-positive V3 neurons (Fig. S6E). In both cases, upregulation was specific to the domain-specific TFs Dbx1 and Nkx2.2, but, consistent with the predictions made by the pseudo-temporal ordering, not seen for the more broadly expressed progenitor TFs Pax6 and Sox2 (Fig. 6F and Fig. S6E).

Based on these observations, we conclude that the transcriptome data allows the domain specific reconstruction of gene expression dynamics underlying neurogenesis. Furthermore, this approach identified neurogenesis as a considerable source of heterogeneity in the expression levels of distinct progenitor markers. Similar approaches may help in the future to pinpoint other sources of gene expression heterogeneity and provide insight into the function of specific genes.

## DISCUSSION

Single cell RNA-seq approaches are revolutionizing the study of cell fate specification and tissue development. The systematic and simultaneous description of gene expression in thousands of cells is accelerating the discovery of new cell types and functions, as well as offering unprecedented insight into the gene regulatory networks that control cell identity (Wagner et al., 2016). In this study we have used scRNA-seq to document the transcriptional signatures of 21465 cells isolated from cervical and thoracic regions of the mouse neural tube during the developmental period that the neuronal subtypes comprising the spinal cord are generated. This sheds light on the changing gene expression profiles that characterise neural tube development and forms the basis of a molecular atlas of the developing mammalian spinal cord.

### An atlas of spinal cord gene expression

The number of cells needed to generate a comprehensive atlas of a tissue depends on multiple factors including the number of cell types and the molecular differences between the cell types (Shekhar et al., 2016). There is a large diversity of cell types in the spinal cord. More than 50 distinct pools of MNs have been documented at limb levels of the spinal cord (Dasen et al., 2005; Landmesser, 2001; Philippidou and Dasen, 2013). Moreover, the combinatorial expression of nineteen TFs have been proposed to generate multiple subpopulations of V1 interneurons (Bikoff et al., 2016; Gabitto et al., 2016; Sweeney et al., 2018). Even though some of this diversity may arise at developmental stages later than those analysed in our study, it is likely that our analysis underestimates the diversity of cell types in the spinal cord. Increasing the number of cells sampled, particularly at e12.5 and e13.5, and improving the sensitivity of the methods to increase the complexity of the analysed transcriptomes might reveal further cell type diversity. In addition, neuronal subtypes vary along the rostral caudal axis of the neural tube (Hayashi et al., 2018; Philippidou and Dasen, 2013; Sweeney et al., 2018), but to ensure the feasibility of sample collection and data analysis, we restricted our analysis to cervical and thoracic regions of the neural tube. Hence, our dataset will not represent cell types unique to lumbar and sacral spinal cord. Nevertheless, despite these limitations, we were able to detect all the progenitor populations and major neuronal subtypes described in cervical and thoracic regions of the spinal cord. Reassuringly, the proportions of the different cell types recovered matched previously determined cell type proportions in the spinal cord (Kicheva et al., 2014), suggesting that the dissection and single cell preparation methods did not substantially bias the composition of the transcriptomes recovered. Consistent with this, relatively small subclasses of several of the neuronal subtypes were readily detected. These included small populations of interneurons such as V3, V2a and dI6 and specific subtypes of MNs including LMC and MMC MNs.

Although much progress has been made to reverse engineer the gene regulatory network responsible for pattern formation and cell fate specification in the neural tube (reviewed in Briscoe and Small, 2015; Lai et al., 2016), it remains incomplete. This study makes available detailed information on the transcriptional state of the entire population of neural tube cells and will likely accelerate efforts to comprehensively map the transcriptional network. To assemble the single-cell transcriptomic atlas, we took advantage of the extensive prior knowledge of gene expression to analyse and organise the transcriptome data (Fig 1D,E). This allowed us to expand the molecular description of cell types and define new subdivisions of neurons. For example, the data revealed that Nr2f1 (also known as COUP-TF1) is enriched in Foxp1 expressing MNs of the lateral motor column (Fig 4C,F) and we detected Pou3f1 in dI1-dI3 neurons, in addition to the described expression in MNs, V2a and V3 interneurons (Dasen et al., 2005; Francius et al., 2013; Rousso et al., 2008) (Fig 3A,E). We found that Nkx6.2 was expressed in a subpopulation of V2a neurons distinct from Shox2 expressing V2a neurons, whereas Pou2f2 was expressed in a group of V3 neurons located lateral to the Olig3 expressing V3 population. This provides a molecular basis for the further partitioning of V2a and V3 neurons into subpopulations as well as implicating these TFs in the generation of specific neuronal subtypes and providing genetic access to investigate their function. Importantly, experimental assays corroborated the transcriptome predictions, suggesting that the atlas is a generally faithful representation of gene expression in the spinal cord. Further mining the dataset will likely implicate additional TFs in the neural tube gene regulatory network and refine the molecular classification of cell types.

Alongside information on the complement of TFs expressed by individual cells, the analysis also revealed the wider molecular repertoire of each cell type. This substantially extends our knowledge of the molecular profile and biological functions of neural tube cells. We illustrated this by demonstrating that the neurotransmitter class of each neuronal subtype, implied by the transcriptome, matches the previously experimentally observed neurotransmitter type and we document the expression of a set of cell adhesion molecules that distinguishes excitatory and inhibitory interneurons (Fig 4A). As anticipated for a complex developing tissue, gene expression profiles change rapidly over time. Cldn3, which encodes a component of tight junctions, is expressed in progenitors of MNs up to e10.5 and then downregulated (Fig 3B-D). The functional significance of this remains to be determined. Considerable additional analysis will be necessary to explore the full potential of the dataset and advance our understanding of the molecular mechanisms of neural development and the allocation of cell type identity. Moreover, as previously discussed (Clevers et al., 2017; Tanay and Regev, 2017), the combination of the dynamic changes in gene expression, the complexity of gene expression, and the experimental noise inherent in single cell transcriptomic techniques emphasizes the need to develop objective criteria to define and delimit cell types.

### Temporal specification of neuronal subtype identity

The data highlighted a previously underappreciated systematic temporal component to the specification neuronal subtype identity in the neural tube. Temporal mechanisms, in which a sequence of TFs is expressed in succession to determine distinct neuronal identities, are well established in cortical neurogenesis and Drosophila neuroblasts (Holguera and Desplan, 2018). In the spinal cord, the birth date of spinal neurons has been shown to correlate with subtype identity, functional properties and settling position in several specific cases (Hayashi et al., 2018; Hollyday and Hamburger, 1977; Sockanathan and Jessell, 1998; Stam et al., 2012; Tripodi et al., 2011). Whether this is a consequence of a global patterning strategy that involves temporally organised changes in gene expression has been unclear. Analysis of the single cell transcriptome dataset revealed a set of gene expression modules activated synchronously at characteristic developmental timepoints in multiple neuronal classes. This raises the possibility of a coordinated temporal transcriptional programme operating in the neural tube that subdivides neuronal classes based on their time of generation. Such a mechanism would operate in parallel to the spatial patterning mechanisms and provide an opportunity to increase the molecular and functional diversity of cell types generated in the neural tube. In addition, coordinating the timing of the generation of sets of neurons might facilitate mechanisms that control the specific settling patterning and connectivity of different neuronal subtypes. Further analysis of the sets of neurons generated at different timepoints will be necessary.

The coordinated induction of gene modules in multiple neuronal subtypes raises might result from a global transcriptional change in the neural progenitors from which they are generated. In the spinal cord, a transcriptional cascade of Sox9 and Nfia/b in progenitors underlies the progressive activation of gliogenesis (Deneen et al. 2006; Kang et al. 2011). Sox9 expression begins between e9.5 and e10.5, while Nfia/b is expressed from e11.5 (Deneen et al. 2006; Kang et al. 2011). However, the forced expression of these TFs does not repress neurogenesis and neural progenitors continue to produce neurons for a considerable period of time after their expression commences (Deneen et al. 2006). The onset of expression of Sox9 coincides with the switch of V1 neurons from Renshaw cells (Onecut expressing) to Foxp2 expressing neurons (Stam et al. 2012; Kang et al. 2011). This raises the possibility that the induction of TFs, such as Sox9 and Nfia/b, changes the competency of neural progenitors to give rise to specific neuronal subtypes and that neural progenitors within a single domain are different over time. In this view, the same mechanism responsible for the activation of gliogenesis in neural progenitors would also serve as a mechanism to generate neuronal diversity in a coordinated manner throughout the spinal cord. Experiments are required to test this hypothesis and to investigate whether similar principles apply to other regions of the nervous system.

Extrinsic signals may also contribute to the temporal stratification of neuronal subtypes. Recent in vitro work suggested that retinoic acid promotes Renshaw cell identity in V1 neurons (Hoang et al. 2018) and the timepoint of their generation correlates with expression of the RA-synthesizing enzyme Aldh1a2 in the adjacent somites (Niederreither et al. 1997). Thus, besides promoting neurogenesis (Novitch et al. 2003), somite-derived RA may be a determinant for early Onecut-positive neuronal subtypes. Another candidate for mediating the temporal progression of neuronal subtypes is TGFβ signalling, which has been shown to suppress early born neuronal identities in favour of later born cell types in multiple regions of the nervous system (Dias et al. 2014). Lastly, we observed the expression of FGF ligands in multiple neuronal subtypes, while neural progenitors at later stages up-regulated FGF receptor3 (Fgfr3) and FGF-binding protein 3 (Fgfbp3) (Gonzalez-Quevedo et al. 2010; Kang et al. 2011). Taken together, multiple secreted signalling molecules may determine into which neuronal subtypes progenitors in the spinal cord differentiate at specific timepoints.

Temporal stratification of neuronal subtypes by shared sets of TFs is probably only one of many mechanisms to generate neuronal diversity in the spinal cord. Recent work suggests the existence of scores of V1 neuron subtypes (Bikoff et al. 2016, Gabitto et al. 2016, Sweeney et al. 2018). This number appears too high for conventional long-range developmental mechanisms. Nevertheless, these neuronal subtypes form in reproducible numbers and occupy stereotypic positions in the spinal cord (Bikoff et al. 2016, Gabitto et al. 2016). One possibility is that the local proximity of multiple neuronal subtypes generates local signalling environments in which neurons experience a combination of multiple signals that expands their diversity. A well-known precedent for such a mechanism is the acquisition of LMCm identity in response to retinoic acid secreted by Aldh1a2+ lateral LMC neurons (Sockanathan and Jessell 1998). In this view, the distribution of neuronal subtypes in the spinal cord is an emergent property in which the spatial and temporal patterns of neurogenesis are intricately linked. The transcriptome resource will provide a framework to analyse this possibility, as it implicates multiple TFs in the establishment of specific interneuron subtypes and can be used to investigate which signalling molecules are produced by different subtypes.

### Dynamics of gene expression during neurogenesis

We have previously demonstrated that upregulation of Olig2 precedes MN formation in vivo and in vitro (Sagner at al. 2018). Pseudotemporal ordering of the progenitor to neuron transition for multiple progenitor domains predicted the transient upregulation of domain specific TFs during neurogenesis (Fig 6C). We confirmed this experimentally for Olig3 in dp2/3, Dbx1 in p0 and Nkx2.2 in p3 progenitors (Fig 6D,E and Fig S6A-D). The success of the pseudotemporal ordering in identifying this behaviour, previously masked in conventional assays by asynchrony of differentiation within a population of progenitors, highlights the resolution afforded by single cell transcriptomes into the sequence of developmental events. Determining the mechanism responsible for the upregulation of TFs during neurogenesis requires further investigation.

We propose that the upregulation of domain-specific TFs prior to neurogenesis might reinforce neuronal specificity during differentiation (Fig. 6G). At early developmental stages, morphogen gradients establish discrete domains of progenitor identities along the DV axis by inducing distinct TFs (Alaynick et al. 2011; Briscoe and Small 2015; Le Dreau and Marti 2012). The TFs form a gene regulatory network that establishes progenitor identities by repressing not only adjacent progenitor identities but also a wide range of alternative fates (Nishi et al. 2015, Kutejova et al. 2016). The dynamics of the gene regulatory network results in progenitors undergoing a succession of changes in TF expression, mediated by repressive interactions, during their specification. The lower levels of domain specific TFs in progenitors, prior to neurogenesis, might therefore represent a mechanism to ensure sensitivity to morphogen inputs and to facilitate cell state transitions in response to morphogen inputs. Such a mechanism, however, could be prone to the generation of mixed neuronal identities as neurogenesis commences (Ericson et al., 1996). The upregulation of the domain specific TFs during neurogenesis might serve to consolidate the appropriate identity and to prevent the initiation of a mixed neuronal identity during differentiation. Thus, the upregulation of domain specific TFs during neuronal differentiation provides a means to enhance the fidelity of spinal cord patterning.

In conclusion, we describe a resource that documents the molecular diversity and cellular composition of the developing mouse neural tube. The data allows the identification of genes and regulatory modules that combinatorially define specific cell types. Moreover, the analysis suggests a temporal axis contributing to neuronal diversity that accompanies the well characterised spatial patterning of neural progenitors. Together, the data provides new opportunities to understand gene function and to genetically target cells for visualization, gene targeting, or manipulation and will support efforts to understand the structure and function of the healthy as well as diseased or damaged spinal cord.

## Supporting information

## ACKNOWLEDGEMENTS

We are grateful to the Advanced Sequencing and Scientific Computing Facilities of the Francis Crick Institute. We acknowledge Monica Tambalo for sharing the protocol for single-cell embryo dissociation for zebrafish embryos that we adapted and optimized for mouse embryos. We thank Bennett Novitch, Thomas Jessell, Susan Morton, Thomas Müller, Carmen Birchmeier and Francois Guillemot for kindly sharing antibodies. A.S. receives funding from an HFSPO postdoctoral fellowship (LTF000401/2014-L), T.R. received funding from an EMBO long term fellowship (ALTF 328-2015). This work was supported by the Francis Crick Institute, which receives its core funding from Cancer Research UK (FC001051), the UK Medical Research Council (FC001051), and Wellcome (FC001051); and by the European Research Council (ERC) under the European Union’s Horizon 2020 research and innovation programme (Grant Number 742138).

## AUTHOR CONTRIBUTIONS

Conceptualization, J.D., T.R., J.B. and A.S.; Methodology, J.D.; Software, J.D.; Formal Analysis, J.D.; Investigation, J.D., T.R. and A.S.; Resources, A.E. and M.M.; Writing – Original Draft, J.D., T.R., J.B. and A.S.; Writing – Review and Editing, J.D., T.R., M.M., J.B. and A.S., Visualization, J.D., T.R. and A.S., Supervision, J.B., Funding Acquisition, T.R., J.B. and A.S.

## MATERIALS AND METHODS

### Animal welfare

Animal experiments in the Briscoe lab were performed under UK Home Office project licenses (PD415DD17) within the conditions of the Animal (Scientific Procedures) Act 1986. Outbred UKCrl:CD1 (ICR) (Charles River) mice were used for this study.

### Immunofluorescent staining and recording of spinal cord sections

Mouse spinal cord tissues were fixed in 4% paraformaldehyde (Thermo Scientific) in PBS, cryoprotected in 15% sucrose, embedded in gelatine, and 14 µm sections taken (Sasai et al. 2014). A complete list of primary antibodies including manufacturer and used dilutions is provided in Table S3. Secondary antibodies used throughout this study were raised in donkey (Life Technologies; Jackson Immunoresearch; Sigma; Biotium). Alexa488 and Alexa568 conjugated antibodies were used at 1:1000 dilution; Alexa647 conjugated antibodies were used at 1:500 and CF405M conjugated secondary antibodies at 1:250.

Images were acquired on a Zeiss Imager.Z2 microscope equipped with an Apotome.2 structured illumination module and a 20x air objective (NA=0.75). 5 phase images were acquired for structured illumination. Z-stacks consisted of 8 sections separated by 1 µm were acquired. Individual optical slices are shown. If required, adjacent images were acquired with 5-10% of overlap and stitching was performed in Fiji using the “Grid/Collection stitching” plugin (Preibisch et al. 2009).

### Sample preparation for single cell RNA sequencing

Outbred CD1 females were timed mated to generate mouse embryos of the specified stages. The morning of the vaginal plug was defined as embryonic stage (e)0.5. For neural tube dissection, cervical and thoracic sections of single mouse embryos were dissected in Hanks Balanced Solution without calcium and magnesium (HBSS, Life Technologies 14185045) supplemented with 5% heat-inactivated fetal bovine serum (FBS). The samples were then incubated on FACS max cell dissociation solution (Amsbio T200100) with 10X Papain (30U/mg Sigma10108014001) for 11min at 37°C to dissociate the cells. To generate a single cell suspension, samples were transferred to HBSS, with 5%FBS, rock inhibitor (Y-27632 10uM, Stem Cell Technologies), and 1X non-essential amino acids (11140035), disaggregated through pipetting, and filtered once through 0.35um filters and once through 0.20um strainers (Miltenyi Biotech 130-101-812). Quality control was assayed by measuring live/cell death, cell size and number of clumps. Samples with a viability above 65% were used for sequencing. 10,000 cells per sample were loaded for sequencing.

### Single cell generation, cDNA synthesis and library construction

A suspension of 10,000 single cells was loaded onto the 10X Genomics Single Cell 3’ Chip, and cDNA synthesis and library construction was performed as per the manufacturer’s protocol for the Chromium Single Cell 3’ v2 protocol (10X Genomics; PN-120233). cDNA amplification involved 12 PCR cycles.

### Nucleic acid sequencing protocol

Libraries for the samples were multiplexed so that the number of reads matched one lane per sample, and sequenced on an Illumina HiSeq 4000 using 100bp paired-end runs. Libraries were generated in independent runs for the different samples and for different time points.

### Alignment and preparation of scRNA-seq data

Demultiplexing, alignment, filtering, as well as barcode and UMI counting were performed using Cell Ranger (version 2.2.0, 10X Genomics, under default settings), which uses STAR aligner. The aggregated gene-barcode matrix was normalized by subsampling reads from higher-depth libraries until all samples had an equal number of confidently mapped reads per cell. Further analysis was performed using R.

### Quality filtering

We excluded cells having more than 6% UMI counts associated with mitochondrial genes and expressing less than 500 genes.

### Data analysis

We partitioned the cells into 13 progenitor and 12 neuronal populations using the following two-step strategy. First, we determined the global cell identities by associating each cell to the closest target population state defined by a list of known marker genes (Figure 1B). The binarized levels were obtained using a threshold of 2 UMI counts and cell-to-target distances were calculated by Euclidean distance. Second, each progenitor and neuronal population was further partitioned using the same approach using the list of marker genes shown on Figure 1D,E.

### Combinatorial testing for differential expression

To classify the whole transcriptome by categories of spatial dorso-ventral patterns, we performed a series of differential expression tests. The analysis was performed independently on progenitors and neurons. In both cases, the procedure was identical: for each gene, 2^N^-2 (resp. N=13 for progenitor domains and N=12 for neuronal domains) approximate χ^2^ likelihood-ratio tests were run between a null hypothesis that models gene expression as a function of a single sample (no predictor) versus an alternative hypothesis modeling gene expression as a function of two samples; the “positive” sample being one of the potential population combinations and the “negative” sample the complementary combination. Hence, all combinations were tested. The tests were run using Monocle (version 2.6.4) *differentialGeneTest* function with gene level distributions modeled as negative binomial distributions with fixed variance (option *expressionFamily=negbinomial*.*size*)), as recommended for UMI datasets). Each gene was then associated with the population combination having obtained the highest likelihood. The gene list was trimmed using significance (P value < 10^-9^) and fold-change (log2-fc > 2) cutoffs. We also ensured that the remaining genes had a minimal average level of expression (0.2 UMI) and a ratio of expressed cells (10% in progenitors, 8% in neurons) in the positive samples.

The gene sets highlighted in Figure 3A were obtained by intersecting the differentially expressed genes with the following GO terms: GO:0098609 for cell-cell adhesion and GO:0006836 for neurotransmitter transport.

### Subclustering of neuronal populations

To investigate the diversity of neuronal subtypes, we independently subdivided the neuronal populations from the mantle zone into 11 dorsal-ventral domains. For each domain, we pooled together the associated neuronal cells and applied the following procedure:

1. Unbiased identification of transcriptomic features

We reasoned that relevant transcriptomic features involve interacting genes demonstrating concerted patterns of expression. From the initial set of ∼5000 expressed genes, we selected the genes that showed highest Spearman correlation with at least three other genes, lowering the correlation cutoff until retaining ∼2000 genes. These genes were then grouped and further filtered with the following 3-step iterative method:

a. The remaining genes were grouped into gene modules by performing a hierarchical clustering using the Spearman dissimilarity matrix of the UMI counts and Ward’s agglomeration criterion. The number of modules were set heuristically to 200 to produce compact and homogeneous groupings.
b. A first filtering criterion was applied to test whether enough cells were expressing the genes comprising the gene module. For each cell, we obtained an average expression level per module by averaging the z-scored log-transformed expression levels of all genes belonging to the module. These gene module averaged levels were then binarized independently by using a parameter-free adaptive thresholding method (R function *binarize.array()* from the ArrayBin package). A cell was considered to express a gene module if the associated Boolean value was true. Modules which were expressed in fewer than five cells were excluded.
c. A second filtering criterion was then applied to test whether cells were expressing the gene module with consistently high levels over most of the genes in the module. We binarized the z-scored log-transformed expression levels of all the remaining genes independently. Then, for each module, we calculated the ratio of Boolean values in cells expressing the module (as defined in b.). We excluded modules where less than 40% of these Boolean values were true.

The iterative loop was terminated when the number of gene modules converged, i.e. when no gene module was excluded in the last iteration. A summary table indicating the number of expressed genes, correlated genes, filtered genes, and gene modules per domain is available in Table S4.

2. Curated selection of the relevant features and neuron type clustering

The gene modules were carefully scrutinized using a list of known neuronal marker genes. We isolated the gene modules related to neuronal identities (Table S2). These identities were obtained by performing a hierarchical clustering of the cells, using Euclidean distances between z-scored log-transformed expression levels of the remaining genes.

### Neuronal trajectories and Pseudotime reconstruction

To project the whole neural dataset into a space that reveals cell state transitions during neurogenesis and progenitor maturation, we aimed at identifying a set of genes with similar dynamics in each dorsal-ventral domain. In order to reduce any bias toward overrepresented populations, we used a resampled dataset containing approximately the same number of cells per timepoint and dorsal-ventral domain. The unbiased gene module identification pipeline described above was used to identify 200 gene modules of concerted patterns of expression in the resampled dataset. Among them, 4 modules were retained as they comprised, respectively, the pan-progenitor marker Sox2, the pan-neuronal marker Tubb3, the early progenitor marker Lin28a and the late progenitor marker Fabp7 (114 genes). An extra step was taken to ensure that no dorsal ventral bias occurred in the remaining genes by testing for differential expression in response to their DV position (Monocle 2.6.4, *differentialGeneTest* function, p-value < 5e-15). 14 genes were excluded and 100 retained (Fig. S6A). After taking the log of the normalized UMI counts (“median ratio” normalization method), Principal Component Analysis was then applied to the 100-dimensional resampled dataset. The resulting PC1-PC2 plane was then populated by multiplying the whole (log-normalized) neural dataset by the eigenvector matrix. Neurogenic pseudotime orderings of each cell were mapped to the PC1 coordinates.

Finally, we reconstructed, independently for each domain, smoothed profiles of gene expression along pseudotime by fitting spline curves (Monocle 2.6.4, *genSmoothCurves* with 3 degree of freedom). Each profile obtained with less than 20 expressed cells was set to zero.

### Data availability

Single cell RNA sequencing data are available via ArrayExpress (http://www.ebi.ac.uk/arrayexpress/experiments/E-MTAB-7320). All other relevant data has been uploaded as Supporting information files. Supplemental tables are available at https://github.com/juliendelile/MouseSpinalCordAtlas.

### Source code availability

The R analysis script developed for this paper is available at https://github.com/juliendelile/MouseSpinalCordAtlas.

## SUPPLEMENTAL FIGURE LEGENDS

**Figure S1:**
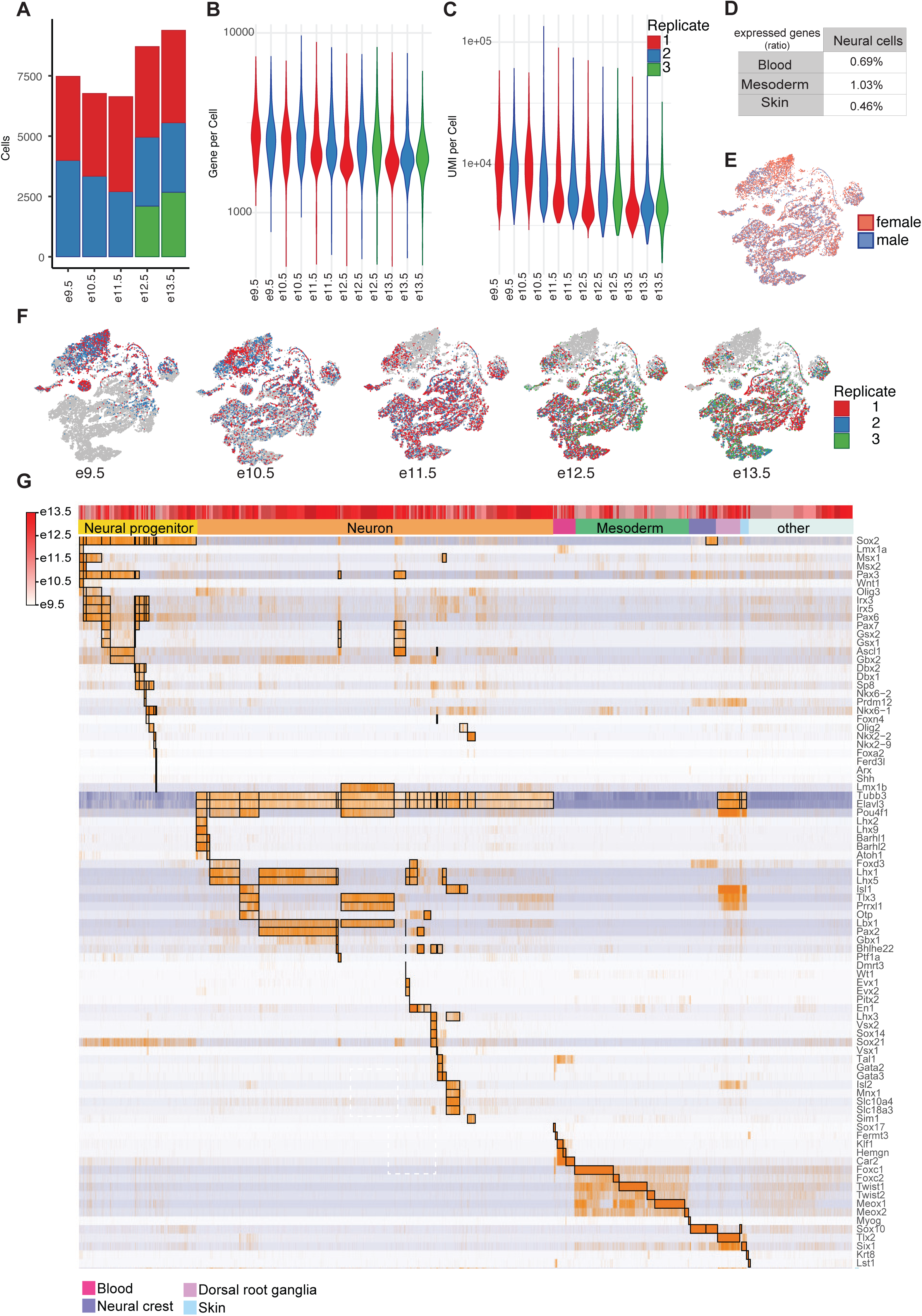
Further characterization of the single cell transcriptome dataset. (A) Number of cells per replicate and timepoint. (B) Number of genes per cell for each replicate. (C) Number of unique molecular identifiers (UMIs) per cell for each replicate. (D) Ratio of blood (Sox17, Fermt3, Klf1, Hemgn, Car2), mesoderm (Foxc1, Foxc2, Twist1, Twist2, Meox1, Meox2) and skin (Krt8) markers expressed in neural cells. (E,F) tSNE-plots showing the distribution of male and female cells in the dataset (E), and the different replicates per timepoint (F). (G) Heatmap showing read distributions of the genes used to assign cell identities to the single cell transcriptomic profiles. Boxed regions indicate the cell identities in which the respective gene is expected to be expressed. Bars at the top of the heatmap indicate sample age (grey early, late red), and cell identity as color coded in Figure 1.

**Figure S2:**
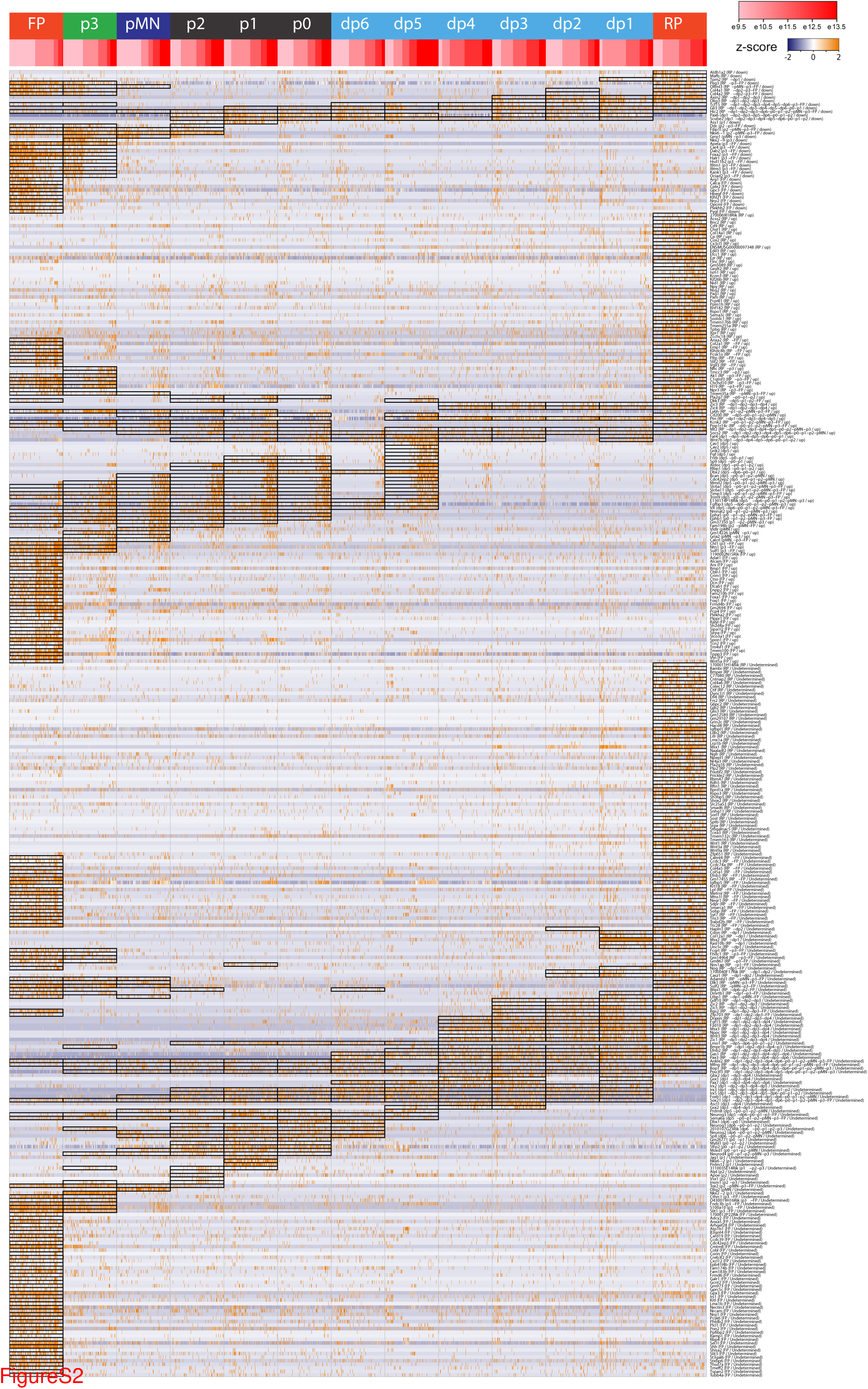
Differentially expressed genes between the different neural progenitor domains. Heatmap showing z-scored UMI count distributions of the genes identified as differentially expressed. Domains predicted to express a gene are indicated by boxes. Cells within each domain are ordered by sample time. Rows are ordered according to the correlation between gene expression and sample time (first downregulated genes, r < -.2, then upregulated genes, r > .2, and finally undetermined trends).

**Figure S3:**
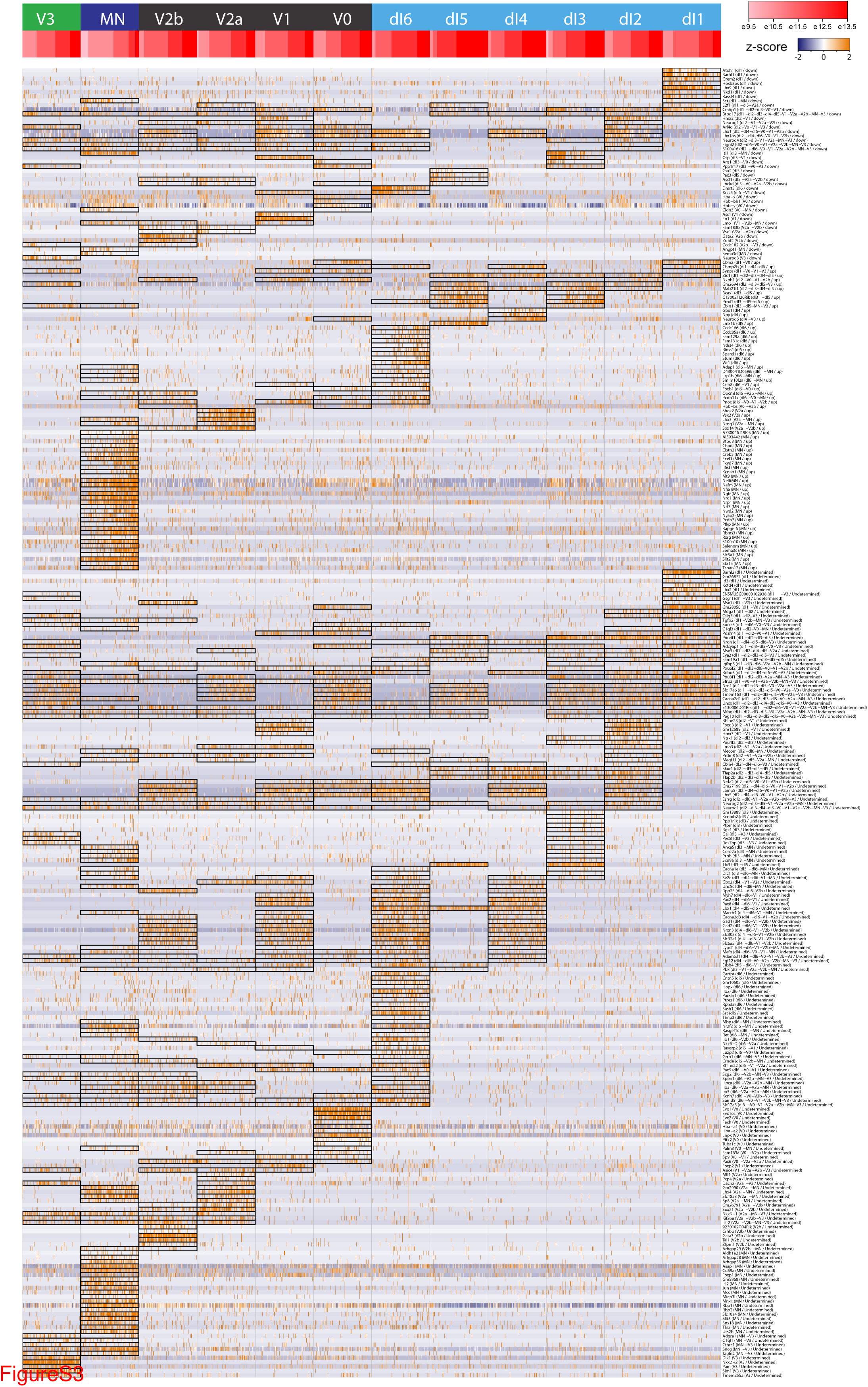
Differentially expressed genes between the different neuronal domains. Heatmap showing z-scored UMI count distributions of the genes identified as differentially expressed. Domains predicted to express a gene are indicated by boxes. Cells within each domain are ordered by sample time. Rows are ordered according to the correlation between gene expression and sample time (first downregulated genes, r < -.2, then upregulated genes, r > .2, and finally undetermined trends).

**Figure S4:**
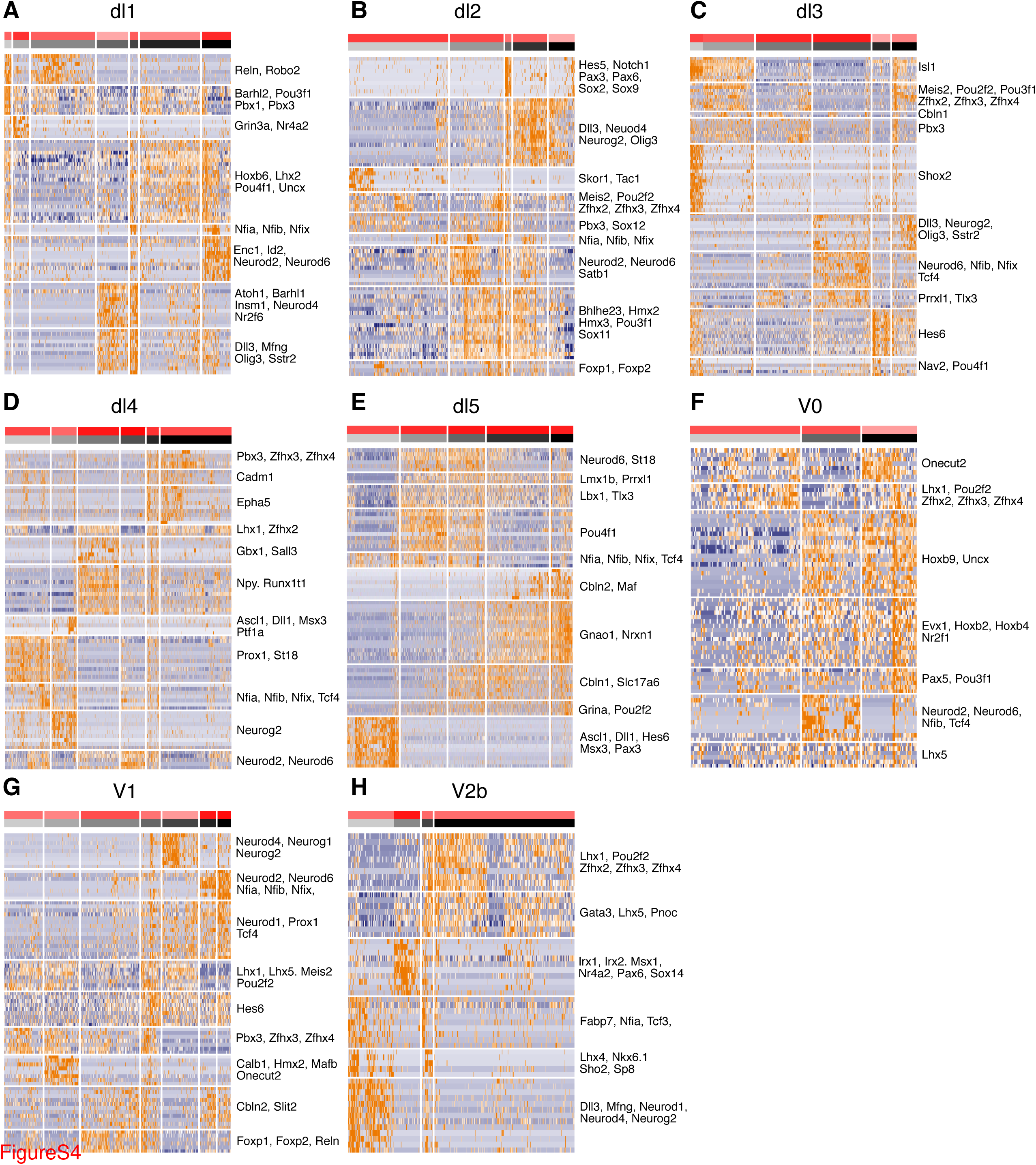
Hierarchical clustering of dI1, dI2, dI3, dI4, dI5, V0, V1 and V2b neurons. Characterization of neuronal subtypes by hierarchical clustering of the indicated domains. Hierarchical clustering was performed using the curated gene modules available in Table S2. A subset of genes included in the modules are indicated on the righthand side.

**Figure S5:**
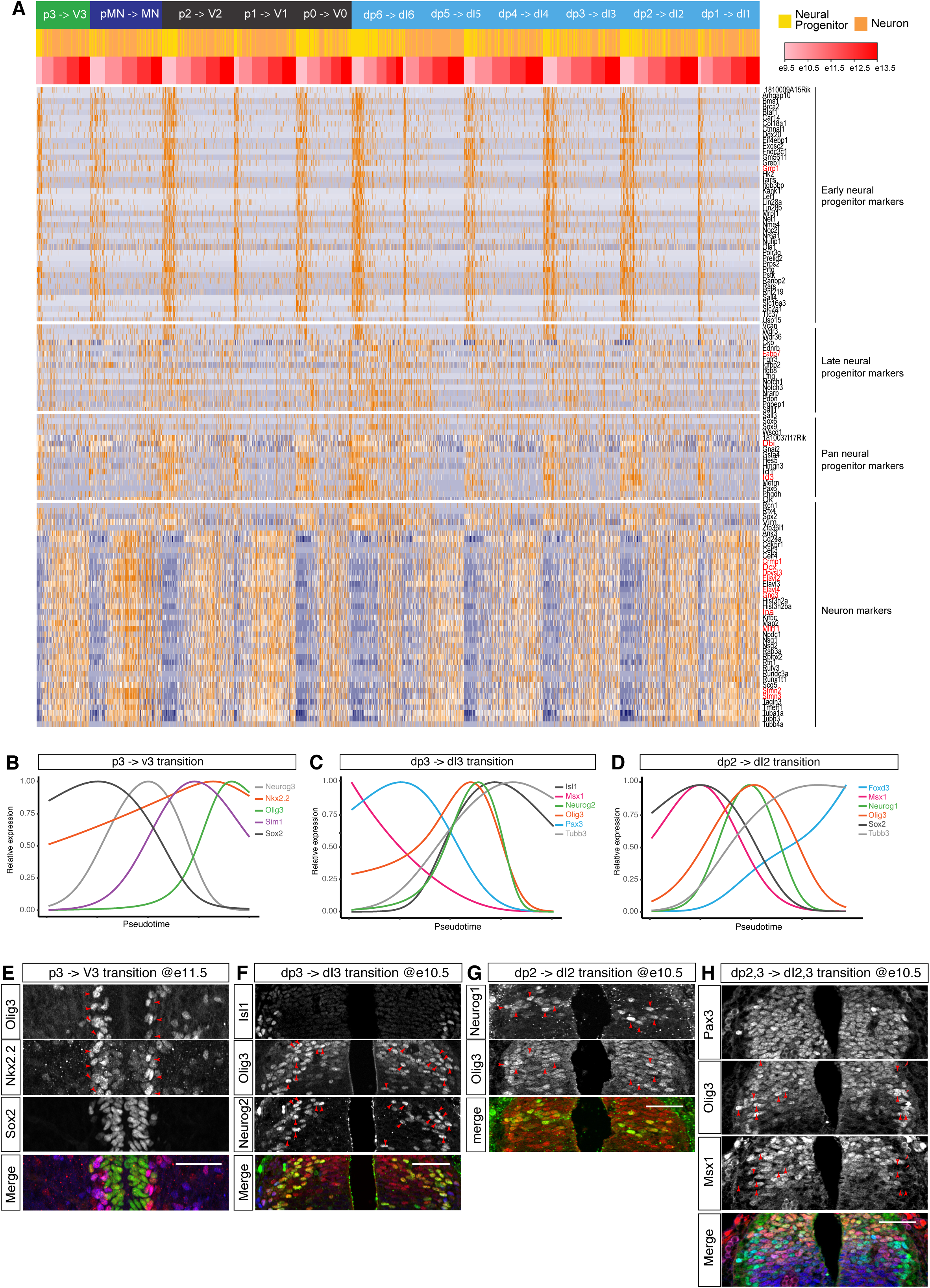
Pseudotime analysis reveals gene expression dynamics underlying neurogenesis in several domains. (A) Heatmap showing the 4 genes modules identified as similarly expressed in all dorso-ventral domains and involved in either neurogenesis and progenitor maturation. Among the 114 genes, red gene labels indicate genes excluded from the following analysis because of significant dorso-ventral bias. (B) Smoothed expression profile of the neurogenic trajectory from p3 to V3 shows increasing levels of Nkx2.2 during neurogenesis. Note that Nkx2.2 in contrast to other neural progenitor markers continues to be expressed in V3 neurons. (C) Smoothed expression profile of the neurogenic trajectory from dp3 to dI3 shows a transient upregulation of Olig3 prior to neurogenesis that coincides with the maximal expression of Neurog2. (D) Smoothed expression profile of the neurogenic trajectory from dp2 to dI2 shows a transient upregulation of Olig3 prior to neurogenesis that coincides with the maximal expression of Neurog1. (E) Upregulation of Nkx2.2 in Olig3+ (red arrows) V3 neurons compared to Sox2+ Nkx2.2 p3 progenitors at e11.5. (F) Upregulation of Olig3 in neurogenic dp3 progenitors labelled by Neurog2 (red arrows). (G) Specific upregulation of Olig3 is anti-correlated with Msx1 (red arrows). The broadly dorsal progenitor marker Pax3 is homogeneously expressed in dp2 and dp3 domains. (H) of Olig3 in neurogenic dp2 progenitors labelled by Neurog1 (red arrows). Scale bars = 50 µm

## SUPPLEMENTAL TABLES

(S1) Knowledge matrix used to identify cell types (see Fig. 1D,E). Columns indicate the cell type classes and rows the genes. A gene contributes to a class definition if the associated value is 1.

(S2) Curated gene lists used to cluster the neuronal subtypes in each dorsal-ventral domain.

(S3) List of primary antibodies including manufacturer and the dilutions used.

(S4) Summary table indicating the number of genes and modules for each step of the gene module identification method used to cluster the neuronal subtypes.

